# Evaluating β-glucanases as cell wall-permeabilising agents against *Phytophthora agathidicida* oospores

**DOI:** 10.64898/2026.05.08.723360

**Authors:** Elise Pierson, Josie C. Mainwaring, Wayne M. Patrick, Monica L. Gerth

**Affiliations:** School of Biological Sciences, Victoria University of Wellington, Wellington 6012, New Zealand

**Keywords:** Oomycetes, glycoside hydrolases, kauri dieback, chemical control, phenol-sulfuric acid assay

## Abstract

The persistence of specialised survival spores produced by microbial pathogens represents a primary bottleneck in the management of plant diseases. In oomycetes, these spores (known as oospores) are largely impervious to chemical control, allowing them to persist in soil and initiate new infection cycles over many years. A prominent example is the soil-borne pathogen *Phytophthora agathidicida*, the causal agent of kauri dieback disease, where long-lived oospores hinder conservation efforts in native forests. The resilience of oospores is attributed to their thick wall composed of complex β-glucan layers that render the oospores impermeable to most conventional biocides. Here we have investigated an enzyme-based approach for weakening the oospore cell wall. We searched enzyme databases to select β-glucanases targeting a variety of linkages found in *Phytophthora* oospore walls. Eight of these β-glucanases were successfully purified and tested for their digestive activity against intact oospores *in vitro* using a phenol-sulfuric acid assay. We showed that combining these enzymes was crucial to achieve significant digestion through synergies and additive effects. The optimal combination, comprising 1,3-, 1,6-, and 1,3(4)-β-glucanases, was evaluated for its ability to permeabilise oospores to five biocides typically effective only on other, more sensitive lifecycle stages of the pathogen. Using a live/dead fluorescence assay, we observed that the effects of the membrane-targeting biocides were potentiated in oospores that were pre-treated with the β-glucanase mixture. Our results highlight enzymatic cell wall permeabilisation as a promising strategy toward improved management of oospore persistence in kauri forest soils and against broader oomycete threats.

**Keypoints:**

- Our phenol-sulfuric acid assay can be used to screen for oospore-degrading enzymes.
- Synergistic enzyme combinations are essential for effective oospore wall digestion.
- Enzyme pre-treatment sensitises oospores to membrane-targeting biocides.

## Introduction

Many microbial plant pathogens produce long-term survival structures to persist in soil or plant debris for years, even under harsh or nutrient-poor conditions (Latijnhouwers et al. 2003; Vega et al. 2019). Among the eukaryotic pathogens, oomycetes (water moulds) produce thick-walled sexual spores known as oospores. These durable structures remain dormant but viable for extended periods, enabling the pathogen to withstand seasonal or chemical stress and then reinitiating infection when conditions become favourable (Judelson 2009). Members of the oomycete genus *Phytophthora* exemplify this strategy, using oospores as survival and dispersal stages in the diseases they cause in crops, forests, and natural ecosystems worldwide. Oospore persistence is one of the greatest obstacles to effective management of *Phytophthora* diseases (Brasier et al. 2022; Erwin and Ribeiro 1996; Hausbeck and Lamour 2004; Judelson and Blanco 2005).

The resilience of oospores can be attributed to their distinctive cell wall architecture. Typically measuring 2.5-3.0 µm thick, the wall comprises several biochemically distinct layers (Beakes et al. 1986; Hegnauer and Hohl 1978). Early studies on *P. cactorum* and *P. megasperma* showed that two insoluble β-glucans constitute about 80 % of the structure. The major component (∼70 %) is a highly branched, noncellulosic β-(1,3)(1,6)-glucan that forms an amorphous inner matrix adjacent to the plasma membrane, while a cellulosic β-1,4-glucan (∼10 %) forms microfibrils concentrated in outer layers and possibly extending inward as a structural backbone (Bartnicki-Garcia and Lippman 1967; Beakes and Bartnicki-Garcia 1989; Lippmann et al. 1974; Mélida et al. 2013; Tokunaga and Bartnicki-Garcia 1971; Wang and Bartnicki-Garcia 1980; Zevenhuizen and Bartnicki-Garcia 1969). This multilayered, β-glucan matrix provides a physical barrier that limits penetration of chemical disinfectants and protects the oospore against environmental stresses (Scott et al. 2009; Stasz and Martin 1988).

Enzymatic degradation of microbial cell walls is a well-established antimicrobial approach, exploited both clinically and agriculturally (Giovannoni et al. 2020; Ullah and Marcianò 2024). For controlling *Phytophthora*, previous efforts have focused on microbial antagonists that secrete cell wall-degrading enzymes (De la Cruz-Quiroz et al. 2018; Martins et al. 2022; Picard et al. 2000; Valois et al. 1996), mulch-linked cellulase activity (Downer et al. 2001; Richter et al. 2011), or purified β-glucanases (Bartnicki-Garcia and Lippman 1967; Kaur et al. 2020). However, these investigations focused on lifecycle stages other than oospores. Early work demonstrated that β-1,3-glucanases could digest isolated oospore wall fragments (Beakes and Bartnicki-Garcia 1989), but the vulnerability of intact oospores to enzymatic degradation remains untested.

We have addressed this knowledge gap for *P. agathidicida*, which causes an incurable root rot disease in the long-lived New Zealand kauri tree (*Agathis australis*) (Bradshaw et al. 2020; Mainwaring et al. 2023). The limited chemical control options for *P. agathidicida* are ineffective at killing oospores (Bellgard et al. 2009; Dick and Kimberly 2013; Lacey et al. 2021; McCarren et al. 2009), and physical methods such as heat-treating soil are impractical beyond greenhouse settings (Bellgard et al. 2018; Horner et al. 2019). Here, we evaluated whether β-glucanase treatment could permeabilise the oospore cell wall. Candidate enzymes were identified through manual database screening, then tested individually and in combination for their ability to release soluble wall carbohydrates from intact oospores. The most active enzyme mixture was then assessed for direct biocidal activity and for its ability to increase oospore permeability to agriculturally relevant chemicals, including Sterigene, copper sulfate, metalaxyl-M, mandipropamid, and nikkomycin Z.

## Materials and Methods

### Materials

Plasmids encoding recombinant β-glucanases were synthesised by GenScript by cloning codon-optimised sequences bearing an N-terminal His_6_-tag followed by a TEV protease cleavage site into pET28a(+) between NcoI and XhoI restriction sites. Chemicals were purchased from Sigma Aldrich unless otherwise specified.

### Enzyme expression and purification

For each β-glucanase, *E.coli* BL21(DE3) GOLD electrocompetent cells were transformed with the expression plasmid and plated on LB (Formedium) agar containing 50 µg mL^-1^ kanamycin (Fisher). Cultures were inoculated from a single colony and grown in LB containing 50 µg mL^-1^ kanamycin at 37 °C and 200 rpm until OD_600nm_ 0.5-0.8. Protein expression was induced with 0.1 or 0.2 mM IPTG (GoldBio) and cultures were further grown for 18 h at 18 °C. Bacteria were harvested by centrifugation, resuspended in a 50 mM HEPES pH 7.5 300 mM NaCl 20 mM imidazole buffer supplemented with 1 mg/mL lysozyme, Viscolase (A&A Biotechnology) and EDTA-free Protease Inhibitor Cocktail (Abcam), and lysed by sonication. The lysate was clarified by centrifugation and purified on a HisTrap FF column (Cytiva) using an AKTA Start FPLC system (Cytiva) and isocratic elution in 50 mM HEPES pH 7.5 300 mM NaCl 500 mM imidazole. The elution buffer was exchanged for storage buffer (50 mM HEPES pH 7.5 150 mM NaCl) using a PD-10 desalting column (Cytiva). Protein concentration was determined spectrophotometrically at 280 nm using the molar extinction coefficient calculated from the sequence using Benchling Biology Software (2025) (https://benchling.com) and purity was assessed by SDS-PAGE. The enzymes were concentrated to around 5 mg/mL by ultrafiltration, aliquoted, snap-frozen in liquid nitrogen, and stored at –80 °C.

### Oospore culture and isolation

*P. agathidicida* isolate NZFS 3770 (Cox et al. 2022) was used in all experiments. Oospores were produced and isolated as previously described (Lacey et al. 2021) with oospore growth medium containing 20% [v/v] clarified V8 broth supplemented with 10 mg/L β-sitosterol and growth times of 8-10 weeks. Oospore purity and numbers were estimated using disposable haemocytometers chips (Incyto) and an inverted microscope. Centrifugation-resuspension steps were repeated to wash the oospores until no mycelial contamination visible under the microscope remained. Purified oospores (50,000-100,000 cells/mL) were stored at 4 °C in the dark.

### Phenol sulfuric method for the determination of β-glucanase oospore digestion activity

Oospore digestion reactions were carried out at 37 °C for 18 h with 2.5 μM of each enzyme and 150,000 oospores per mL in reaction buffer (100 mM sodium citrate pH 6.0, 1 mM MgCl_2_.6H_2_O). The reactions were stopped by pelleting the oospores at 16,000 × g for 5 min and transferring the supernatant to a fresh tube. The supernatant was mixed with an equivalent volume of 5% [w/v] phenol. The resulting solution was vigorously mixed with 95-98% sulfuric acid in a 1:5 ratio and incubated at room temperature for 60 min. Enzyme activity was measured as a function of absorbance at 485 nm, which was correlated to the mass concentration of total soluble carbohydrate released via a D-glucose calibration curve. Blank reactions were prepared identically to each enzyme treatment but without oospores (enzyme(s) + buffer only). For the untreated control, blanks contained buffer only.

Final data points were obtained by subtracting, from each measurement, the mean concentration of the corresponding blank (triplicate) for that condition.

### Oospore viability assays

The enzyme cocktail was prepared by mixing equimolar amounts of enzymes E1, E2, E3, E4 and X1. Biocide compounds were solubilised at 10 mg mL^-1^ in 1% [v/v] DMSO. To reduce variability due to incubation times and mechanical stress between controls, single treatments and double treatments, treatments were carried out in parallel in two sequential phases for all samples. In the first phase, oospores (∼2000) were incubated with either the enzyme cocktail (for *enzyme+biocide* treatments) or an equal volume of storage buffer (50 mM HEPES pH 7 7.5 150 mM NaCl; for *biocide* treatments, and for buffer, DMSO and autoclaved controls) in 100 µL digestion buffer (100 mM sodium citrate pH 6.0 1 mM MgCl_2_.6H_2_O). The final concentration of each enzyme in the reaction was 2.5 µM. Reactions were incubated in the dark at 22 °C for 18 hours. For the autoclaved control, oospores from the original stock were autoclaved (121 °C 30 min 15 psi) prior to the set up. After incubation, oospores were washed twice by centrifuging at 1,200 × g for 5 min, discarding 90% of the supernatant and replacing it with sterile ultra-pure water. In the second phase, the oospores were incubated at 22 °C for 18 h in the dark with either 100 µg mL^-1^ nikkomycin Z, copper sulfate, metalaxyl-M (Toronto Research Chemicals), or mandipropamid, or 2% [v/v] Sterigene (Oragene) (for *biocide* and *enzyme+biocide* treatments), or with sterile ultra-pure water (for buffer and autoclaved controls), or with 1% DMSO (for DMSO control). Oospores were then washed twice as described above. Treated oospores, as well as an untreated sample from the original oospore stock (for exposure standardisation), were pelleted at 1,200 × g for 5 min and the supernatant was discarded. Oospores were then resuspended in 10 µL fluorescein diacetate at 100 µM and incubated at 37 °C for 20 h in the dark. DiTO-3 (AAT Bioquest) was then added to a final concentration of 6.67 µM, after dilution of the commercial solution 1:5 in DMSO. Oospores were further incubated at 37 °C for 4 h in the dark. Oospores were then washed once to remove excess dye and reduce background fluorescence. Oospore were imaged by microscopy and red and green fluorescence were measured as described in Fairhurst et al. (2023). The untreated and autoclaved samples were used as dye controls and to standardise the exposure times for each channel in each experiment. Five independent experiments were performed in total (n=5). An average of 100 oospores was visually assessed per sample. Images taken in the brightfield, green and red channels were merged using ImageJ (Schneider et al. 2012) and the number of green, red, dual-stained (yellow), and unstained oospores were manually counted.

### Statistical analysis

All statistical analyses were performed using GraphPad Prism 10.4.2. All data is expressed as mean ± SD (standard deviation). Differences between groups were compared using one-way ANOVA followed by Dunnet’s post-hoc test for pairwise comparisons for the phenol sulfuric assay data (α = 0.05), and repeated measures one-way ANOVA with the Geisser-Greenhouse correction followed by Šidák test for the viability assay data (α = 0.05). Effect sizes were estimated using Cohen’s *d* for paired samples, calculated as the mean of the differences (treatment-reference) divided by the SD of the differences for each comparison.

### Accession numbers for strains and amino acid sequences

*P. agathidicida* isolate 3770 (GenBank Assembly: GCA_025722995.1, ICMP 17027), enzyme E1 (GenBank: ABD82280.1), enzyme X1 (GenBank: QPA36577.1), enzyme E2 (GenBank: ABD82251.1), enzyme E3 (GenBank: AAO18342.1), enzyme E4 (GenBank: AXR85444.1), enzyme E5 (GenBank: AAB38548.1), enzyme X2 (NCBI Reference Sequence: WP_011051143.1), enzyme X3 (GenBank: ABX43721.1), *Streptomyces coelicolor* cellulose 1,4-beta-cellobiosidase (NCBI Reference Sequence: WP_011031000.1).

## Results

### Identification and purification of candidate enzymes for digesting oospores

To identify candidate enzymes for oospore cell wall digestion, we searched the BRENDA (Hauenstein et al. 2025) and the Carbohydrate Active Enzymes (CAZy, http://www.cazy.org/ (Drula et al. 2021)) databases for glycoside hydrolases (Enzyme Commission (EC) number 3.2.1.-) specifically active towards β-glucans. We focused our search on 1,4-, 1,3-, and 1,6-β-glucanases targeting the β-1,4- and β-(1,3)(1,6)-glycosidic bonds thought to be found in *P. agathidicida* oospores cell wall and included β-glucanases targeting ‘mixed-linkage’ β-1,3(4)-glucans. We examined the associated literature for bacterial β-glucanases with a pH optimum between 4.5 and 6.5 and retaining at least 50% of their maximal specific activity across temperatures of 20 to 30 °C. These criteria ensured that enzymes would be compatible with field soil pH values (4.5-6.0) and optimum growth temperatures for kauri (23-26°C) (Bieleski 1959). Our search yielded hits from 11 EC classes, with 2 to 8 candidate enzymes per class. The selection was further refined to one enzyme per EC class, based on sequence availability, and maximised activity at 20-30 °C. Two classes whose activities were redundant were eliminated. The final selection comprised nine β-glucanases with complementary cleavage patterns (endo- or exo-acting) and linkage position specificities (**Table 1**). A more detailed version of this table containing kinetic parameters, GH families, products and preferred substrates can be found in Supplementary Information (**Table S1**). All enzymes were successfully expressed in *E. coli* and purified by affinity chromatography, except the cellulose exo-β–1,4–glucobiosidase SCO6546 (EC 3.2.1.91) which could not be purified.

**Table 1.** β-glucanases selected for oospore cell wall digestion.

| Code<sup>a</sup> | Activity [EC 3.2.1.n] | Source, enzyme name | Target linkage(s) | pH opt., other pH (rel. activity) | T opt., lower T (rel. activity) | Ref. |
| --- | --- | --- | --- | --- | --- | --- |
| **E1** | endo-1,3-β-glucanase [39]<br>+ endo-1,6- $\beta$ -glucanase [75] | *Saccharophagus degradans* 2-40<sup>T</sup>, Gly5M | β-1,3<br>+ β-1,6 | 6.0 | **40°C**<br>20°C (80%) | (Wang et al. 2016) |
| **X1** | exo-1,3- β-glucosidase [58] | Moose rumen microbiome, Exg5 | β-1,3 | 5.0-6.0 | **40°C**<br>30°C (90%)<br>20°C (80%) | (Kalyani et al. 2021) |
| **E2** | endo-1,6- β-glucanase [75] | *Saccharophagus degradans* 2-40, Gly30B | β-1,6 | 7.0<br>6.0 (85%) | **40°C**<br>30°C (75%)<br>20°C (65%) | (Wang et al. 2017) |
| **E3<sup>b</sup>** | endo-1,3(4)- $\beta$ glucanase [6] | *Zobellia galactanivorans*, EngA (res. 56-385) | β-1,3<br>+ β-1,4 | 6.0-6.5 | **45°C**<br>25°C (60%)<br>15°C (45%) | (Dorival et al. 2018) |
| **E4<sup>b</sup>** | 1,3;1,4- β-glucan endohydrolase / licheninase [73] | *Bacillus* sp. SJ-10, β-1,3-1,4-glucanase | β-1,4 | 6.0 | **50°C**<br>30°C (80%)<br>20°C (75%) | (Kim et al. 2013) |
| **E5** | endo-1,4- β-D-glucanase / cellulase [4] | *Fibrobacter succinogenes*, CelG | β-1,4 | 5.5 | **25°C** | (Iyo and Forsberg 1996) |
| **X2** | exo- β-1,4-glucanase / exocellulase [74]<br>+ exo-1,3- $\beta$ -glucosidase [58] | *Xanthomonas citri* pv. *Citri*, Bgl3B | β-1,4<br>+ β-1,3 | 6.0 | **35°C**<br>20°C (80%)<br>15°C (60%) | (Vieira et al. 2021) |
| **X3** | cellulose 1,4- $\beta$ -cellobiosidase (reducing end) [176] | *Clostridium phytofermentans*, Cel48 | β-1,4 | 5.0-5.5 | **50-55°C**<br>35°C (65%) | (Zhang et al. 2010) |
| - | cellulose 1,4-beta-cellobiosidase (non-reducing end) [91] | *Streptomyces coelicolor* A(3), SCO6546 | β-1,4 | 5.0 | **50°C**<br>30°C (70%) | (Lee et al. 2018) |

<sup>a</sup> E = endo-acting enzyme, X = exo-acting enzyme
<sup>b</sup> These enzymes act on mixed-linkage glucans only, not on pure 1,3- or 1,4-glucans

### Individual β-glucanases show limited oospore digestion ability

We evaluated the eight purified β-glucanases for their ability to hydrolyse *P. agathidicida* oospores and release soluble carbohydrates in solution, as measured by the phenol sulfuric acid method. Intact, purified oospores were incubated with individual enzymes, after which undigested oospores were separated from the solution and total carbohydrate content of the supernatant was quantified. The results are shown in **Fig. 1.** Only enzyme E5 showed significantly elevated total carbohydrate release (3.9 ± 1.4 µg mL^-1^) above the untreated baseline (0.9 ± 0.3 µg mL^-1^, *p* < 0.01). Enzyme E1 showed modest but non-statistically significant activity (2.5 ± 1.6 µg mL^-1^) while the remaining six enzymes produced no detectable digestion above the untreated baseline.

**Fig. 1.**
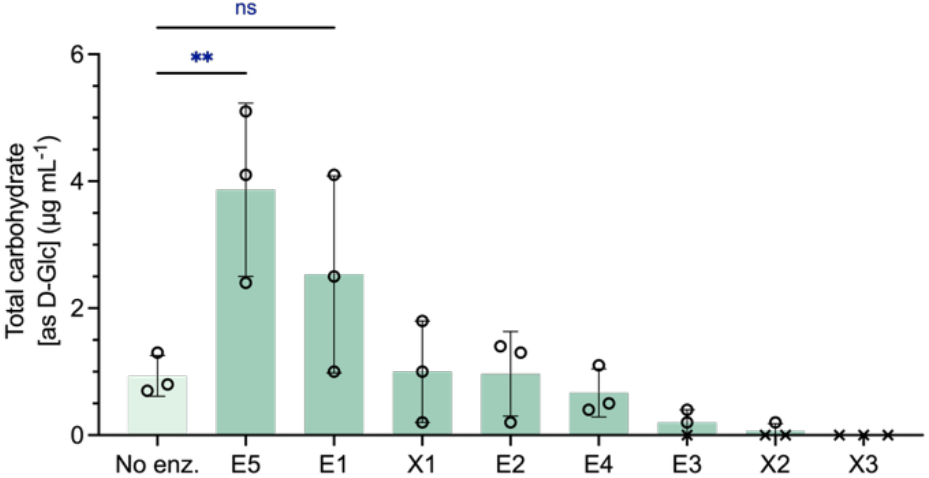
Total carbohydrate released from intact *P. agathidicida* oospores treated with individual β-glucanases, as determined by the phenol sulfuric acid method. No enz. = baseline from untreated oospores. The bar height represents the mean of three biological replicates (n=3, depicted as ○) measured in triplicate. Measurements falling below the limit of detection were set to zero and are depicted as ×. Error bars represent SD. Statistical differences were assessed using one-way ANOVA followed by Dunnet’s post-hoc test for pairwise comparisons. Asterisks indicate significant differences between conditions (** *p* < 0.01)

### β-glucanase cocktails reveal synergies and incompatibilities in oospore digestion

We then combined the enzymes into three cocktails, each tailored to target a type of glucan in the oospore cell wall: β-(1,3)(1,6)-glucans, β-1,4-glucans, and (suspected) β-1,3(4)-glucans (**Fig. 2**). The (1,3)(1,6)-β-glucanase cocktail (E1 + X1 + E2) produced a signal significantly above baseline (6.0 ± 0.7 µg mL^-^1, *p* < 0.001) and on average 2.3× greater than the apparent sum of individual activities (2.6 ± 1.9 µg mL^-1^, *p* < 0.05) indicating a potential synergy (**Fig. 2a**). Similarly, the 1,3(4)-β-glucanase cocktail (E3 + E4) showed clear synergistic effects (**Fig. 2b**), releasing on average 8× more product than the apparent sum released by E3 and E4 individually (5.3 ± 0.3 vs. 0.7 ± 0.5 µg mL^-1^, *p* < 0.0001). In contrast, the 1,4-β-glucanase cocktail (E5 + X2 + X3) had an antagonistic effect (**Fig. 2c**); it released on average 3× less soluble carbohydrates than the total from E5, X2 and X3 individually (1.3 ± 0.3 vs. 3.9 ± 1.4 µg mL^-1^, *p* < 0.05). The activity of the 1,4-β-glucanase cocktail was not significantly different from the baseline.

**Fig. 2.**
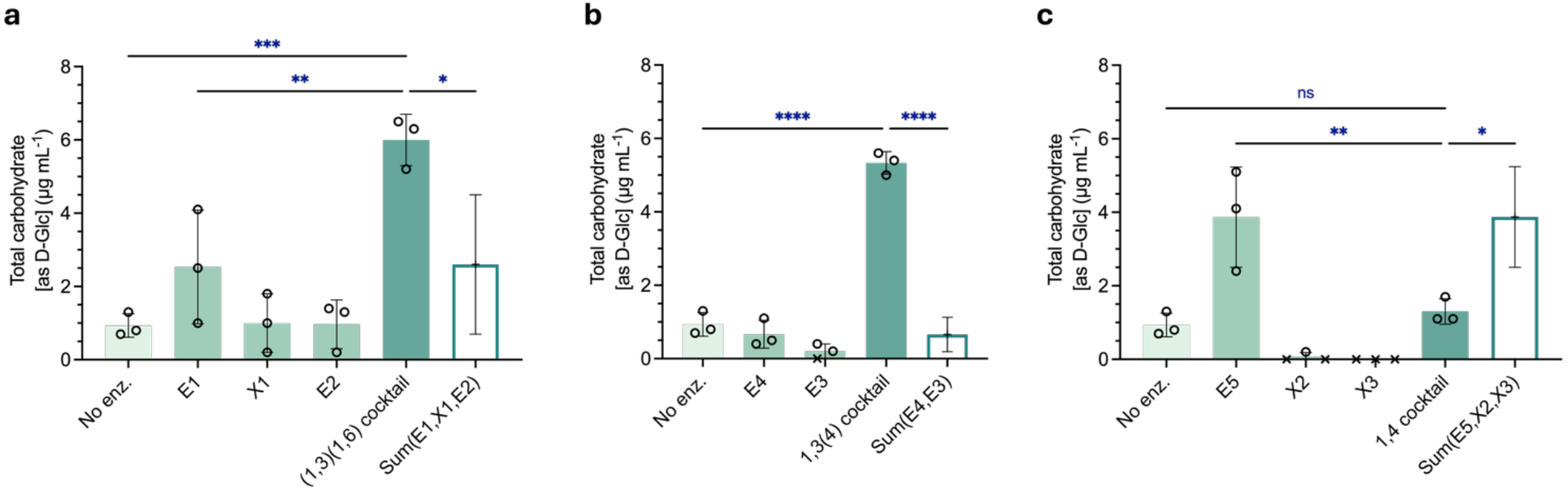
Total carbohydrate released from intact *P. agathidicida* oospores treated with cocktails of β-glucanases and their individual components, as determined by the phenol sulfuric acid method. **a)** cocktail of 1,3- and 1,6-β-glucanases **b)** cocktail of 1,3(4)-β-glucanases **c)** cocktail of 1,4-β-glucanases. No enz. = untreated oospores baseline. The bar height represents the mean of three biological replicates (n=3, depicted as ○) measured in triplicate. Measurements falling below the limit of detection were set to zero and are depicted as ×. Error bars represent SD. White bars depict the arithmetic sum of the signals from individual cocktail components. Statistical differences were assessed using one-way ANOVA followed by Dunnet’s post-hoc test for pairwise comparisons. Asterisks indicate significant differences between conditions (\**p* < 0.05, ** *p* < 0.01, \*\*\**p* < 0.001, **** *p* < 0.0001)

Lastly, the three cocktails were mixed in all permutations to identify the most efficient combination of substrate specificities (**Fig. 3**). The (1,3)(1,6)-β-glucanase and 1,3(4)-β-glucanase cocktails were seen to work in an additive fashion (**Fig. 3a**) and their combination yielded the highest oospore digestion activity (11 ± 1.0 µg mL^-1^), significantly exceeding the activity of the cocktail containing all 8 enzymes and those of the other pairwise combinations (**Fig. 3b**). As expected from the incompatibility seen between enzymes E5, X2, and X3 (**Fig. 2c**), all the combinations containing the 1,4-β-glucanase cocktail exerted antagonistic effects when mixed with the other cocktails (**Fig. S1**).

**Fig. 3.**
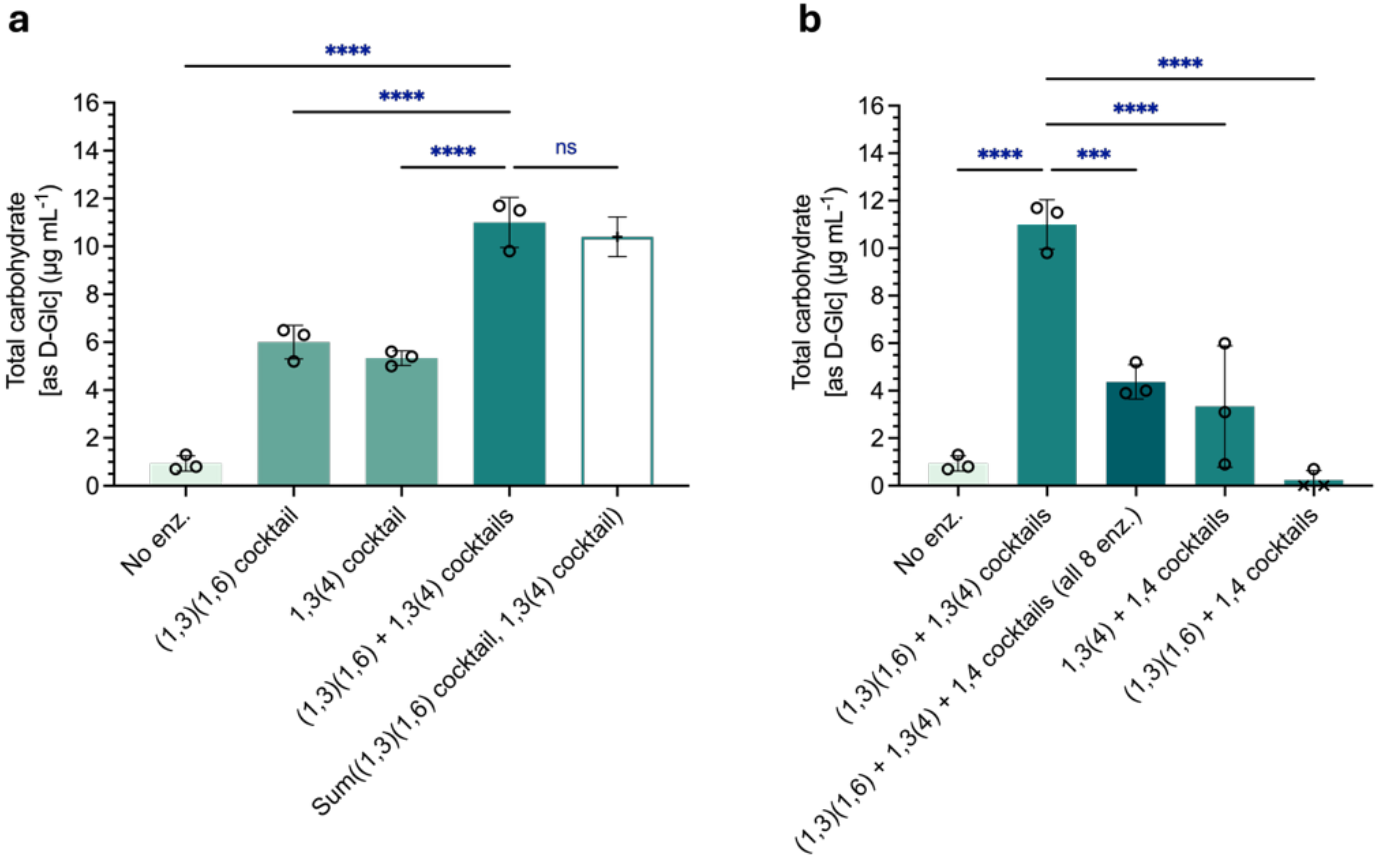
Total carbohydrate released from intact *P. agathidicida* oospores treated with combinations of β-glucanases cocktails, as determined by the phenol sulfuric acid method. **a)** Combination of (1,3)(1,6)- and 1,3(4) β-glucanase cocktails **b)** Pairwise combinations of (1,3)(1,6)-, 1,3(4)- and 1,4-β-glucanase cocktails and full mixture thereof. No enz. = untreated oospores baseline. The bar height represents the mean of three biological replicates (n=3, depicted as ○) measured in triplicate. Measurements falling below the limit of detection were set to zero and are depicted as ×. Error bars represent SD. The white bar depicts the arithmetic sum of the signals from individual combination components. Statistical differences were assessed using one-way ANOVA followed by Dunnet’s post-hoc test for pairwise comparisons. Asterisks indicate significant differences between conditions (\**p* < 0.05, ** *p* < 0.01, *** *p* < 0.001, **** *p* < 0.0001)

### β-glucanase treatment potentiates the effect of biocides on oospore viability

We tested whether the combination of (1,3)(1,6)-β-glucanase and 1,3(4)-β-glucanase cocktails directly affected oospore viability, or increased oospore susceptibility to the biocides Sterigene, copper sulfate, mandipropamid, metalaxyl-M, and nikkomycin Z. Treated oospores were stained with a mix of two fluorescent dyes:

1. a green dye (fluorescein diacetate) marking metabolically active cells with intact membranes, and
2. a red dye (DiTO-3) marking non-viable cells with disrupted membranes. For each treatment, we microscopically enumerated the proportions of green (intact membrane, viable cell), red (ruptured membrane, dead cell), dual-stained (compromised membrane, dying cell), and unstained oospores (**Fig. S2**). Red and dual-stained oospores (i.e. all oospores stained with DiTO-3) were combined into a single “damaged” category, as our primary interest was to quantify structural damage, reflected by increased numbers of both dead and compromised oospores. In parallel, we were also interested in loss of viable cells, reflected by decreased numbers of green-only oospores.

To evaluate the effect of *biocide-only* (or *enzymes-only*) treatments, we compared the percentages of damaged oospores and viable oospores between the treated samples and their respective baseline controls (buffer or buffer + 1% DMSO). *Enzyme+biocide* combinations were then compared with the corresponding *biocide-only* treatments to evaluate any potentiating effect of the enzyme pre-treatment. Changes in viability and damage were statistically non-significant for all comparisons. This was unsurprising given the inherent metabolic and genetic heterogeneities of oospore populations (Boutet et al. 2010; Vercauteren et al. 2011), which limit statistical power for precise quantification. To distinguish true null effects from potentially meaningful biological activity masked by high variability and limited replication (n=5), we calculated effect sizes (Cohen’s *d*) from paired differences. Cohen’s *d* measures the difference between control and treatment groups in standard deviation units, with thresholds defining “small” (0.2) “medium” (0.5), “large” (0.8), and “very large” (1.2) effects (Cohen 1988; Sawilowsky 2009). Reporting effect sizes alongside statistical significance has been encouraged to interpret findings in light of both chance and magnitude (Ialongo 2016; Nakagawa and Foster 2004). Results are shown in **Table 2**. A detailed version of this table, including statistical analyses, is provided in **Supplementary Table S2**.

**Table 2.**
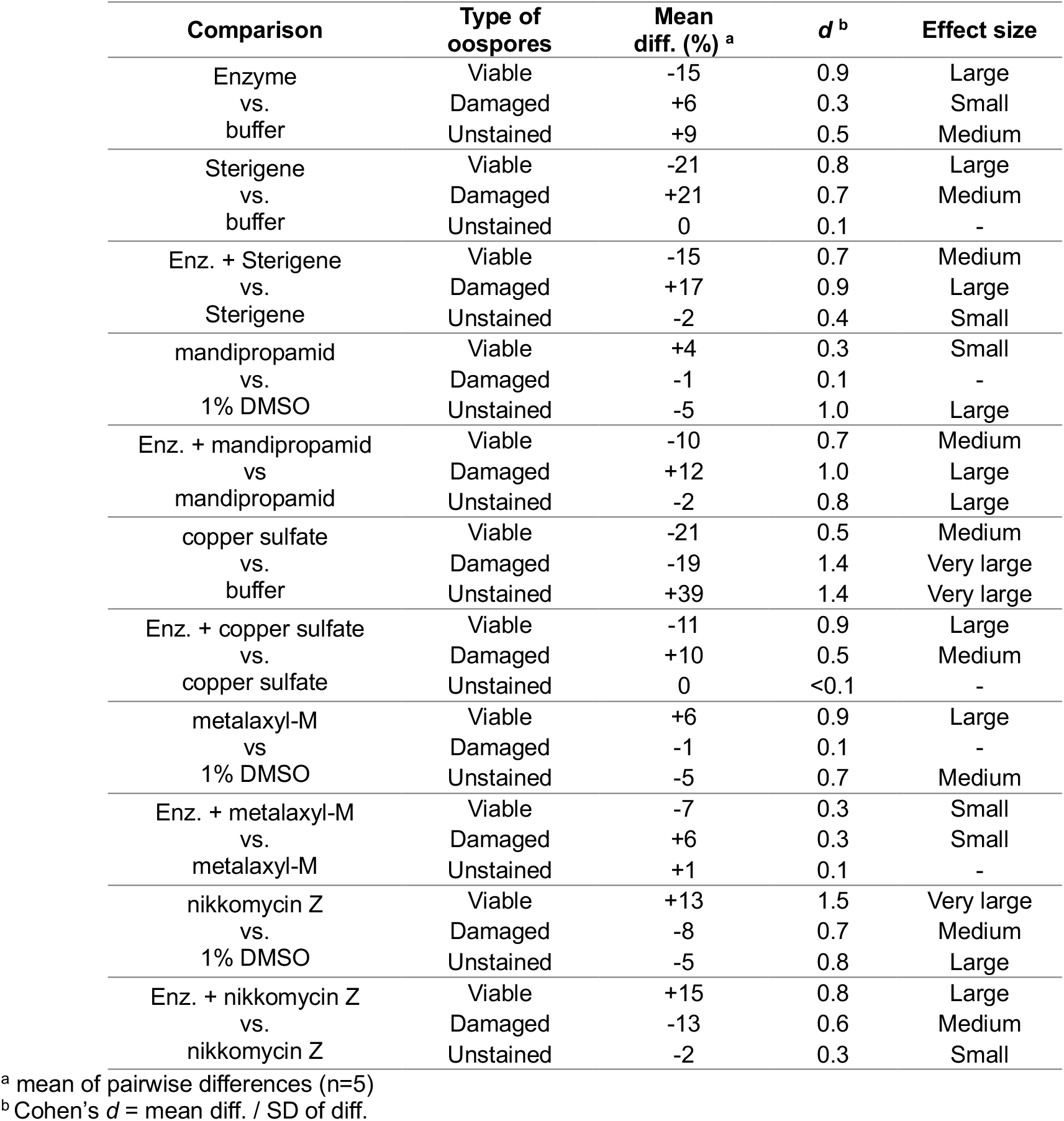
Changes in viable (green-stained), damaged (red- and dual-stained) and unstained oospores in treated oospore samples as determined by live/dead fluorescent staining using fluorescein diacetate/DiTO-3 dyes and microscopic enumeration, and their relative effect sizes estimated as Cohen’s *d*.

Four treatments decreased the proportion of viable (green-stained) oospores and increased the proportions of damaged (red- and dual-stained) oospores with medium-to-large effect sizes, suggesting biocidal trends (**Table 2, Fig. 4**). Among these, *Sterigene-only* treatment reduced viability (−21%) and increased damage (+21%) compared to buffer, and *enzymes+Sterigene* further reduced viability (−15%) and increased damage (+17%) relative to *Sterigene-only* treated oospores (**Fig. 4a**).

**Fig. 4.**
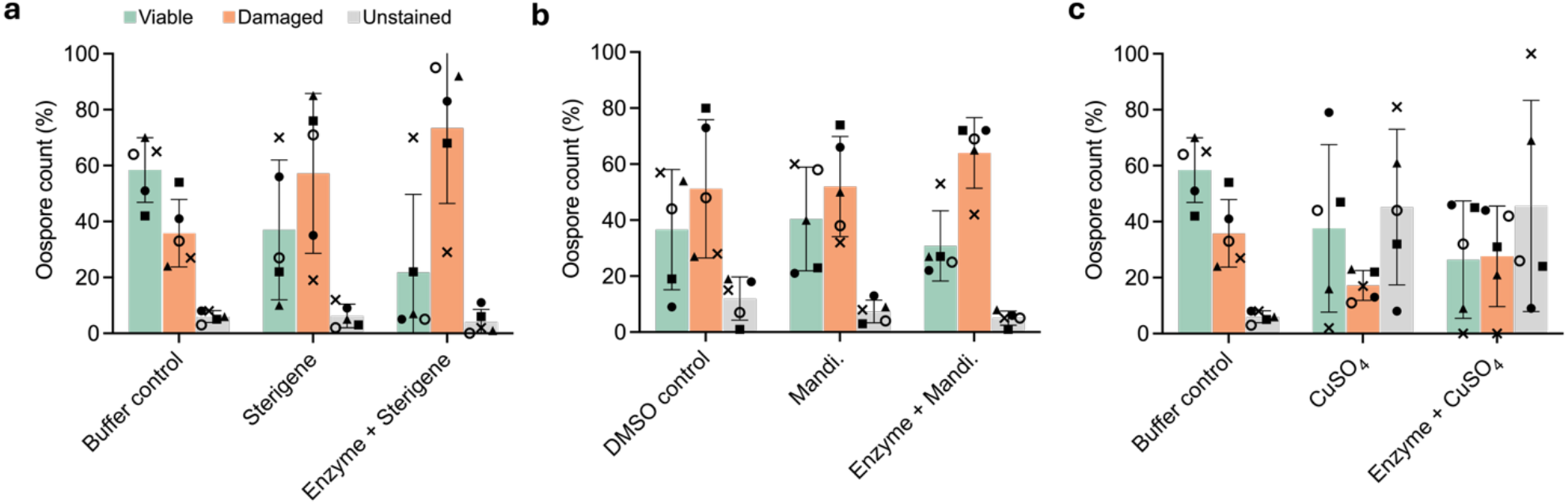
Percentage of green-stained (viable), red- + dual-stained (damaged) and unstained oospores in treated oospore samples as determined by live/dead fluorescent staining using fluorescein diacetate/DiTO-3 dyes and microscopic enumeration. **a)** Oospores treated with buffer (baseline), 2% [v/v] Sterigene, and sequentially treated with the β-glucanase cocktail and 2% [v/v] Sterigene **b**) Oospores treated with buffer + 1% DMSO (baseline), 100 µg mL^-1^ mandipropamid, and sequentially treated with the β-glucanase cocktail and 100 µg mL^-1^ mandipropamid **c)** Oospores treated with buffer (baseline), 100 µg mL^-1^ copper sulfate, and sequentially treated with the β-glucanase cocktail and 100 µg mL^-1^ copper sulfate. The bar height represents the mean percentage of five biological replicates (n=5), each represented by a different symbol (×,▴,•,▪,∘). Error bars represent SD.

*Enzymes+ mandipropamid* also further reduced viability (−10%) and increased damage (+12%) compared to *mandipropamid-only* treatment (**Fig. 4b**), and *enzymes +copper sulphate* reduced viability (−11%) and increased damage (+10%) compared to *copper sulfate-only* (**Fig.4c**).

Interestingly, treatments including nikkomycin Z showed an opposing trend (**Fig. 5a**). With medium-to-very large effect sizes, *nikkomycin Z-only* treatment increased the count of viable oospores (green-stained, +13%) and decreased the count of damaged (red and dual-stained, -8%) oospores compared to the DMSO baseline. *Enzymes+nikkomycin Z* treatment further increased viability (+15%) and decreased damage (−13%) relative to *nikkomycin Z-only* treated oospores.

**Fig. 5.**
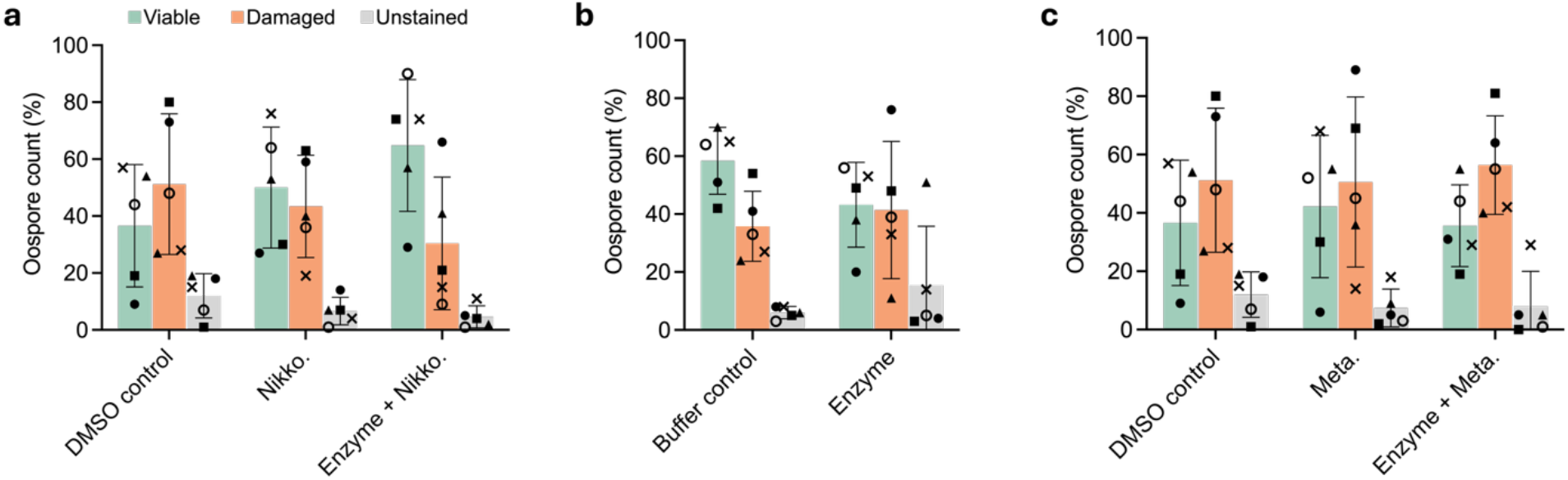
Percentage of green-stained (viable), red- + dual-stained (damaged) and unstained oospores in treated oospore samples as determined by live/dead fluorescent staining using fluorescein diacetate/DiTO-3 dyes and microscopic enumeration. **a)** Oospores treated with buffer + 1% DMSO (baseline), 100 µg mL^-1^ nikkomycin Z, and sequentially treated with a β-glucanase cocktail and 100 µg mL^-1^ nikkomycin Z **b)** Oospores treated with buffer (baseline), and with the β-glucanase cocktail (enzyme) **c)** Oospores treated with buffer + 1% DMSO (baseline), 100 µg mL^-1^ metalaxyl-M, and sequentially treated with a β-glucanase cocktail and 100 µg mL^-1^ metalaxyl-M. The bar height represents the mean percentage of five biological replicates (n=5), each represented by a different symbol (×,▴,•,▪,∘). Error bars represent SD

Some treatments exhibited medium-to-very large effect sizes for increases in the unstained fraction, making the effect of these treatments ambiguous. This is the case for the *enzymes-only* treatment (**Fig.5b**), where the large drop in green-stained oospores (−15%) could be correlated to the increase in unstained oospores (+9%) rather than to a true loss of viability. This is consistent with the only small effect size increase in the damaged fraction. The proportion of unstained oospores also dramatically increased (+39%) in the *copper sulfate-only* treatment (**Fig. 4c**), reflecting drops in both green-stained (−21%) and damaged cells (−19%). Due to its oxidative nature, copper could be interacting with staining mechanisms. Conversely, *mandipropamid-only* (**Fig. 4b**) and *metalaxyl-only* (**Fig. 5c**) treatments showed medium-to-large decreases in the unstained fraction (−5%). These drops correlated with increases in the proportion of green-stained oospores, as there was no change in the damaged fractions. The apparent effects likely reflect increased staining variability in the DMSO control rather than a true increase in viability, suggesting these treatments did not substantially affect oospores.

Finally, *enzymes+metalaxyl* (**Fig. 5c**) showed small-to-negligible effect sizes for changes in viability and damage, indicating likely null effects. The effects of all the treatments are summarised in **Table 3**.

**Table 3.**
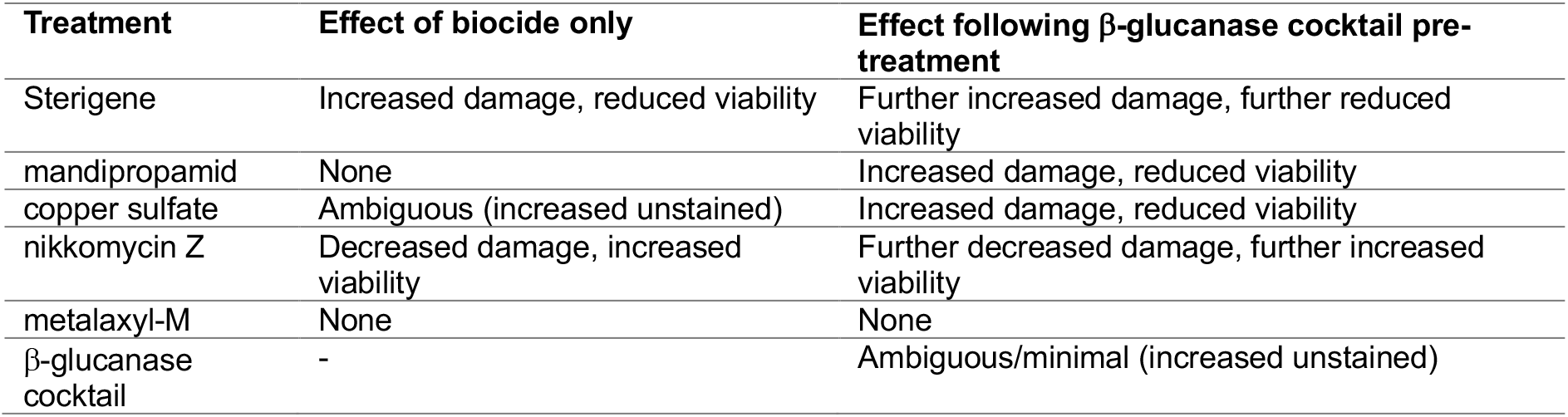
Summary of the effects of biocide treatments, with and without enzyme pre-treatment, on the fractions of viable and damaged oospores as determined by live/dead fluorescent staining using fluorescein diacetate/DiTO-3 dyes and microscopic enumeration

## Discussion

This study provides proof of concept that the walls of intact *P. agathidicida* oospores can be enzymatically modified using selected β-glucanases and that this can alter how oospores respond to chemical treatments. This is significant because oospores represent one of the most recalcitrant *Phytophthora* life stages and demonstrating that their defensive ‘shield’ can be functionally compromised is a critical step toward improving disease management.

Using a phenol–sulfuric assay, we showed that recombinant β-glucanases can hydrolyse cell wall polysaccharides of intact oospores. The most efficient β-glucanase mixture in our experiment digested approximately 8% of cell wall β-glucans by mass. This was calculated using established parameters in which cell wall biomass represents 50% of oospore dry weight and is composed of 80% glucan (Lippmann et al. 1974), with an estimated lyophilised oospore mass of 2.31 ng (Ruben 1975). Our experiment yielded a lower percentage than the 24–34% digestion reported by Beakes and Bartnicki-Garcia (1989) for treatment of isolated cell wall fragments. The discrepancy likely stems from the challenges associated with digesting intact oospores instead of these fragments. Our findings also demonstrate that the phenol–sulfuric assay, applied here for the first time to intact oospores, is a straightforward approach for screening cell wall-degrading enzymes in a biologically relevant context.

From all the enzyme combinations screened, the optimal cocktail contained enzymes that act on β-1,3-glucans, β-1,6-glucans, and mixed linkage β-1,3(4)-glucans. This confirms established knowledge of oospore cell wall composition while providing fresh evidence that extends previous structural models. The efficacy of the mixture containing enzymes acting on β-1,3-glucans and β-1,6-glucans (*i.e.* the (1,3)(1,6)-β-glucanase cocktail containing E1, X1, and E2; **Fig. 2a**) aligns with the fact that branched β-(1,3)(1,6)-glucans comprises ∼70% of the oospore wall (Lippmann et al. 1974).

Unexpectedly, the 1,3(4)-β-glucanase cocktail (*i.e.* E3 and E4) proved nearly as effective, releasing comparable amounts of soluble carbohydrate as the (1,3)(1,6)-β-glucanase cocktail (**Fig. 2b**) and showing additive effects when both cocktails were combined (**Fig. 3a**). This was surprising because the enzymes in the 1,3(4)-β-glucanase cocktail only target chains containing both β-1,3 and β-1,4 linkages (Dorival et al. 2018; Kim et al. 2013). Our results therefore imply the presence of glucans with this mix of linkages in the oospore wall. Such linkages have never been documented in oospore walls, although earlier studies detected them in hyphal walls of *P. cinnamomi* (Zevenhuizen and Bartnicki-Garcia 1969) and more recently in those of oomycetes of the Saprolegniales orders (Mélida et al. 2013). We propose there are crosslinks between the β-1,4-linked cellulosic backbone and the β-1,3-linked amorphous matrix in the oospore wall. This is consistent with a model in which a network of cellulosic microfibrils extends through the amorphous β-1,3-linked layers (Beakes and Bartnicki-Garcia 1989).

Our results illustrate that significant digestion requires a combination of enzymes with complementary linkage specificities and mechanisms of action (*i.e.* both endo- and exo-acting). This has also been observed when enzymatically digesting complex plant cell wall polysaccharides (Khamassi and Dumon 2023; Van Dyk and Pletschke 2012). Oospore cell wall β-glucans are no exception to this principle of synergistic cooperation. However, enzyme compatibility is not universal, and antagonistic effects can arise when the binding of one enzyme physically hinders the access or action of another (Medve et al. 1998; Thoresen et al. 2021). In this study, exo-acting β-1,4-glucanases (X2 and X3) exhibited such antagonism when combined with other enzymes (**Fig. 3b**). We postulate this may result from their binding to the cellulosic glucans concentrated in the outer wall layers, thereby restricting the penetration of matrix-targeting hydrolases.

The optimal enzyme cocktail (refined to exclude the antagonistic exo-β-1,4-glucanases) was subsequently paired with biocides known to be active against the mycelial and zoospore lifecycle stages of *Phytophthora* (Bellgard et al. 2009; Garbelotto et al. 2009; Hinkel and Ospina-Giraldo 2017; Lawrence et al. 2017; Matheron and Porchas 2000; Qu et al. 2016; Wang et al. 2013). In general, enzymatic pre-treatment enabled an effect on oospore viability or enhanced the effect of the biocide alone (**Table 3**). While the nature of the effects differed across different biocides, they are nevertheless all consistent with a model in which the enzymes permeabilize the cell wall and increase access of exogenous molecules (both biocides and fluorescent reporter dyes) to the underlying plasma membrane. For instance, the increased efficacy of the cellulose synthase inhibitor mandipropamid (**Fig. 4b**) suggests that enzymatic weakening of the cell wall allows the biocide to reach its membrane-bound target. Once at the membrane, mandipropamid can inhibit synthesis of the cellulosic backbone and contribute to osmotic lysis of the oospore (Kawai et al. 2023). As defects in cell walls modify permeability to dyes (Jones et al. 2016), enzyme treatment may also facilitate DiTO-3 diffusion into cells whose membranes have already been compromised by the biocide alone. This phenomenon may explain the effects seen with copper sulfate. Copper ions diffuse through cell walls and denature membranes, enzymes and nucleic acids through oxidative stress (Gaetke et al. 2014; Sarkar et al. 2023). In our assay, this mode of action translated into an increase in unstained oospores, likely due to copper both denaturing the intracellular esterases required for fluorescein diacetate hydrolysis and disrupting DNA intercalation by DiTO-3. At the same time, enzyme treatment increased the proportion of damaged oospores by 10% without affecting the unstained fraction (**Table 2**), which suggest enhanced DiTO-3 uptake rather than increased copper penetration.

Finally, the observations for metalaxyl-M and nikkomycin Z provide a check against the possibility that the enzyme treatment itself was toxic, while supporting our model. The lack of effect with metalaxyl-M is consistent with the metabolic dormancy of oospores. This biocide inhibits RNA polymerase I (Chen et al. 2018), and so would lack an active target inside dormant oospores, regardless of wall permeability. Although counterintuitive, the increase in viable (green) staining with nikkomycin Z (**Fig. 5a**) is also consistent with enzyme treatment increasing the permeability to biocides and dyes. More significantly, it also aligns with growing evidence that *Phytophthora* walls contain polymers of chitin (*N*-acetylglucosamine), which historically was not thought to be the case (Mélida et al. 2013). Ullah et al. (2026) recently demonstrated that nikkomycin Z inhibits the chitin synthase of *P. infestans* (*Pi*Chs). *Pi*Chs synthesizes short, soluble chito-oligosaccharides rather than the insoluble chitin found in other cell walls (Ullah et al. 2026). Such chito-oligosaccharides are thought to contribute to cell wall scaffolding in oomycetes (Badreddine et al. 2008). The genome of *P. agathidicida* (Cox et al. 2022) contains a putative chitin synthase gene (locus KNV87_015747), encoding a protein with 94% identity to *Pi*Chs. Moreover, treatment with a chitinase is required to effect protoplast formation in *P. agathidicida* (Hayhurst et al. 2025). We therefore hypothesise that nikkomycin Z affects the integrity of the *P. agathidicida* oospore wall by inhibiting the chitin synthase activity required for chito-oligosaccharide production. However, because *N*-acetylglucosamine is a minor component of oomycete walls (Mélida et al. 2013), the defect appears insufficient to trigger osmotic lysis and consequent red dye (DiTO-3) penetration. Increased wall permeability from this defect could, nevertheless, enhance uptake of the membrane-permeable fluorescein diacetate, leading to increased viable (green) staining. Enzyme treatment appears to accentuate this effect by enhancing access of nikkomycin Z to the membrane-bound chitin synthases.

Collectively, the patterns seen across the biocides demonstrate that targeted enzymatic digestion can bypass the oospore’s physical defences to expose internal vulnerabilities, while also emphasizing that both the biocide’s mode of action and the molecular mechanisms of the stains used *in vitro* are critical considerations when interpreting laboratory results for deployment in the field.

Overall, these findings support a model in which the oospore wall is chemically complex and functionally protective, but not invulnerable. Future work should focus on refining enzyme compatibility, broadening the range of wall-targeting activities (such as investigating chitinase enzymes), and pairing wall digestion with orthogonal methods for evaluating long-term viability. Ultimately, this work identifies cell wall permeabilization as a promising strategy for overcoming one of the most persistent barriers in *Phytophthora* management.

## Statements and declarations

### Author Contribution Statement

MG conceived the general project idea and secured funding for the work. MG, WP, and EP designed this study. EP performed all experiments. EP and JM developed the methodology. EP, WP, and MG analysed the data. EP wrote the first draft of the manuscript; MG and WP contributed to editing and writing, and JM provided comments. All authors reviewed and approved the final version.

