## Supplemental Info for "Evaluating β-glucanases as cell wall-permeabilising agents against *Phytophthora agathidicida* oospores"

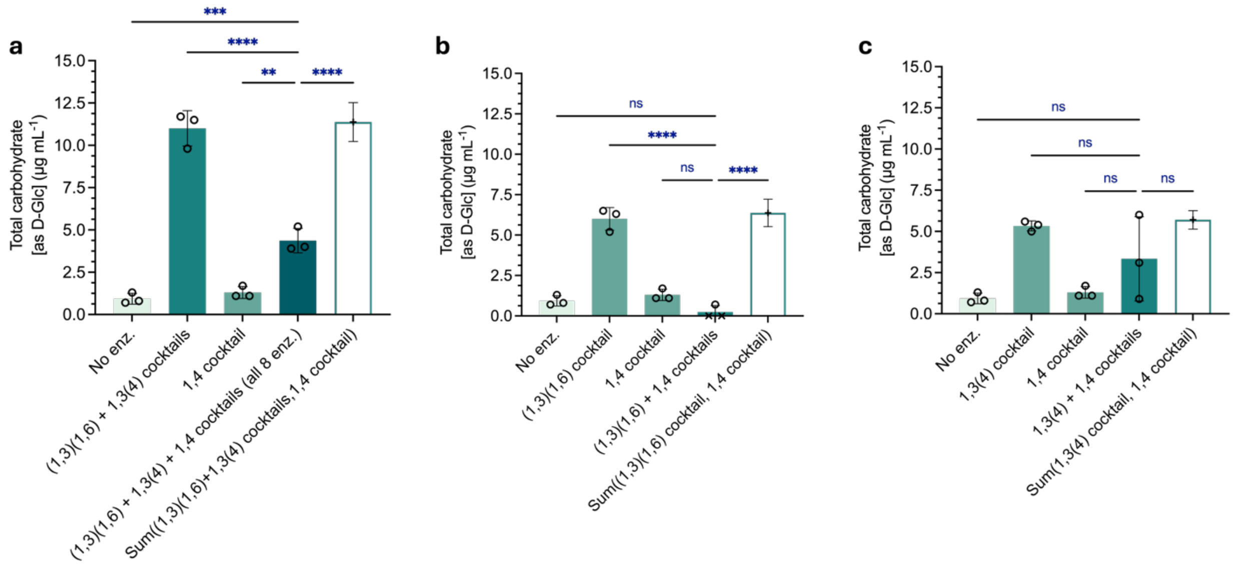


**Fig. S1** Total carbohydrate released in the supernatant of solutions of intact *P. agathidicida* oospores treated with combinations of β-glucanases cocktails and their individual components, as determined by the phenol sulfuric acid method: effect of mixing the 1,4-β-glucanase cocktail with **a)** the combination of **(**1,3)(1,6)-, 1,3(4)- and 1,4-β-glucanase cocktails **b)** the (1,3)(1,6)- β-glucanase cocktail **c)** the 1,3(4)- β-glucanase cocktail. No enz. = untreated oospores baseline. The bar height represents the mean of three biological replicates (n=3, depicted as ○) measured in triplicate. Measurements falling below the limit of detection were set to zero and are depicted as ×. Error bars represent SD. The white bar depicts the arithmetic sum of the signals from individual combination components. Statistical differences were assessed using one-way ANOVA followed by Dunnet's post-hoc test for pairwise comparisons. Asterisks indicate significant differences between conditions (*$p < 0.05$, ** $p < 0.01$, *** $p < 0.001$, **** $p < 0.0001$)

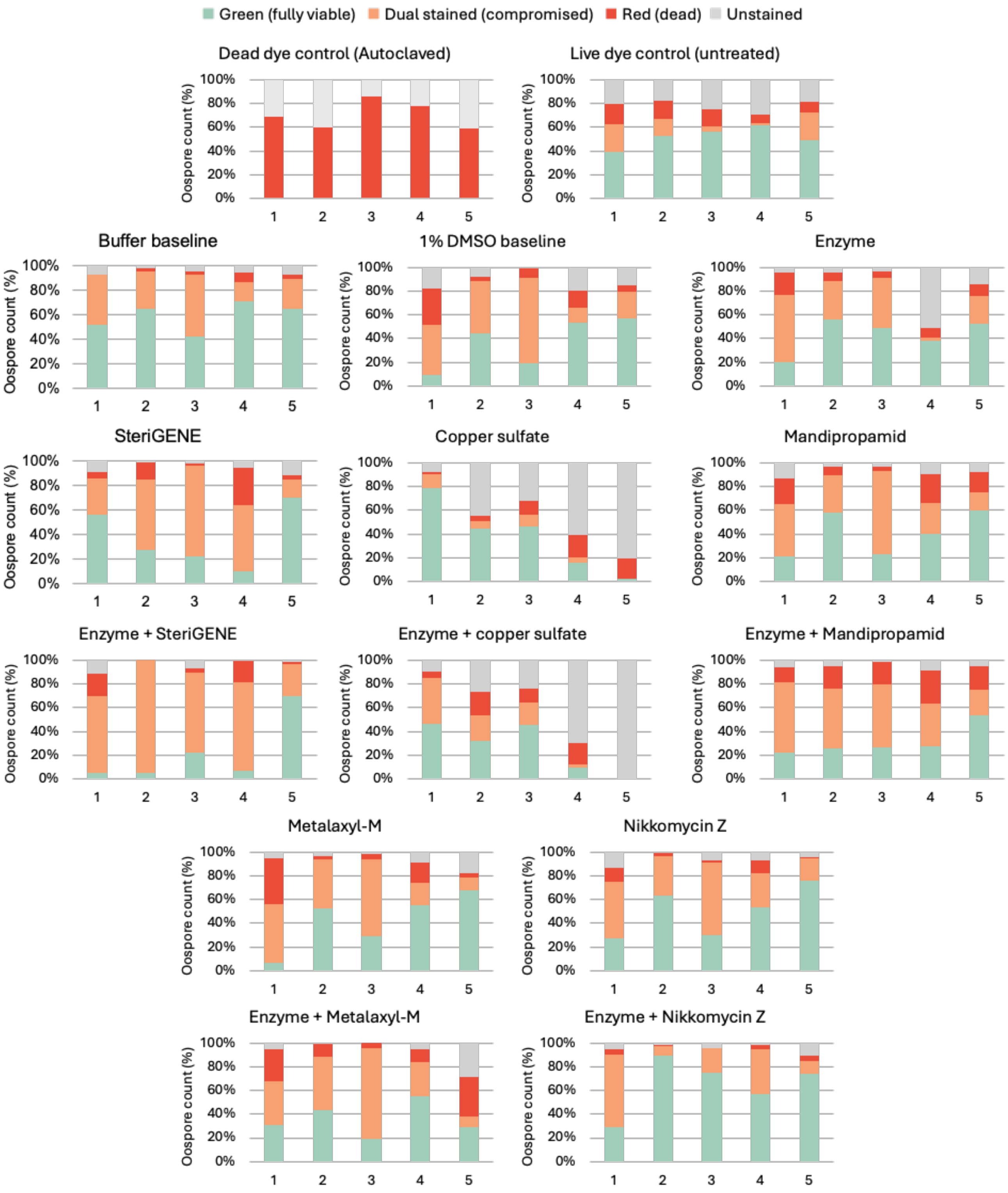


**Fig. S2** Percentage counts of green (fully viable), dual-stained (compromised), red (dead) and unstained oospores as determined by live/dead fluorescent staining using fluorescein diacetate and DiTO-3 dyes and microscopic enumeration. Oospores samples were either autoclaved (positive control for damage), treated with buffer or buffer + 1% DMSO (baseline controls), treated with β-glucanase cocktail (enzyme) only, treated with biocide only (2% [v/v] SteriGENE or 100 μg $mL^{-1}$ biocide), or sequentially treated with the β-glucanase cocktail and the biocide (enzyme + biocide). Number 1 to 5 identify biological replicates.

**Table S1** β-glucanases selected for oospore cell wall digestion and their documented characteristics

| | **Activity (EC 3.2.1.-)** | **Enzyme** | **GH fam.** | **Target linkages** | **Notes** | **Exo/endo** | **Product** | **pH opt.,** other pH (rel. activity) | **T opt.,** lower T (rel. activity) | **Preferred substrate(s),** other substrates (rel. activity) | **$k_{cat}$** | **$K_M$** | **Ref.** |
|---|---|---|---|---|---|---|---|---|---|---|---|---|---|
| **E1** | endo-1,3-β-glucanase (39)<br>+ endo-1,6- β -glucanase (75) | *Saccharophagus degradans 2-40$^T$* Gly5M | 5 | β-1,3<br><br>+ β-1,6 | Very limited action on substrates containing both β-1,3 and β-1,4 bonds | Endo | glucan | 6.0 | **40°C**<br>20°C (80%) | **Laminarin**<br>Pustulan (55.6%) | 16.8 $s^{-1}$<br>- | 10.4 g/L<br>- | (Wang et al., 2016) |
| **X1** | exo-1,3- β -glucosidase (58) | *moose rumen microbiome* Exg5 | 5_44 | β-1,3 | - | Exo (non-red. end) | α-glucose | 5.0-6.0 | **40°C**<br>30°C (90%)<br>20°C (80%) | **L5 β-1,3 oligosaccharide**<br>L4, L3, L2 " (81.7-35.7%)<br>$3^2$-β-D-glucosyl-cellobiose (74.3%)<br>----<br>**pNP-β-D-glucopyranoside**<br>---<br>**Laminarin** | 125.4 $s^{-1}$<br>-<br><br><br>131.1 $s^{-1}$<br><br>122.4 $s^{-1}$ | 1.7 mM<br>-<br><br><br>0.26 mM<br><br>3.72 mg $ml^{-1}$ | (Kalyani et al., 2021) |
| **E2** | endo-1,6- β -glucanase (75) | *Saccharophagus degradans 2-40* Gly30B | 30 | β-1,6 | - | Endo | glucan | 7.0<br>6.0 (85%) | **40°C**<br>30°C (75%)<br>20°C (65%) | **Pustulan**<br>Laminarin (22.4%) | 135.6 $s^{-1}$<br>28.9 $s^{-1}$ | 24.2 g/L<br>100.8 g/L | (Wang et al., 2017) |
| **E3** | endo-1,3(4)- β -glucanase (6) | *Zobellia galactanivorans* EngA (catalytic module: residues 56-385) | 5 | β-1,3 + β-1,4 | Glucose residue whose reducing group is involved in the linkage to be hydrolysed must be substituted at C3. *Zg*EngA preferably cleaves β-1,3 flanked by β-1,4 linkages. | Endo | glucan | 6.0-6.5 | **45°C**<br>25°C (60%)<br>15°C (45%) | **Mixed-linkage glucan**<br>Glucomannan (60.8%)<br>Lichenan (29.4%)<br>Xyloglucan (5.9%)<br>CMC (3.9%) | n.d. | n.d. | (Dorival et al., 2018) |
| **E4** | 1,3;1,4- β-glucan endohydrolase / licheninase (73) | *Bacillus sp. SJ-10* Beta-1,3-1,4-glucanase | 16 | β-1,4 | Not active on pure β-1,3- or β-1,4-glucans | Endo | glucan | 6.0 | **50°C**<br>30°C (80%)<br>20°C (75%) | **Barley β-glucan**<br>Oat β-glucan (4.6%)<br>β-1,3-glucan (2.3%)<br>laminarin (0.7%) | n.d. | n.d. | (Kim et al., 2013) |
| **E5** | endo-1,4- β -D-glucanase / cellulase (4) | *Fibrobacter succinogenes* CelG | 5_4 | β-1,4 | Also hydrolyses β -1,4 linkages in substrates containing both β-1,3 and β-1,4 bonds. | Endo | glucan | 5.5 | **25°C** | **Barley β-glucan**<br>Lichenan (30%)<br>Soluble oats spelt xylan (27%) | n.d. | n.d. | (Iyo & Forsberg, 1996) |
| **X2** | exo- β-1,4-glucanase / exocellulase (74)<br>+ exo-1,3- β -glucosidase (58) | *Xanthomonas citri pv. Citri* Bgl3B | 3 | β-1,4<br><br>+ β-1,3 | - | Exo | glucose | 6.0 | **35°C**<br>20°C (80%)<br>15°C (60%) | **pNP-β-D-glucopyranoside**<br>pNP-β-D-cellobioside (4%)<br>---<br>**β-1,4-glucobiose**<br>Laminarin (20%) | 32.83 $s^{-1}$<br>-<br><br>8.16 $s^{-1}$<br>2.47 $s^{-1}$ | 3.68 mM<br>-<br><br>1.35 mg/mL<br>5.93 mg/mL | (Vieira et al., 2021) |
| **X3** | cellulose 1,4- β -cellobiosidase (reducing end) (176) | *Clostridium phytofermentans* Cel48 | 48 | β-1,4 | - | Exo | cellobiose | 5.0-5.5 | **50-55°C**<br>35°C (65%) | **Regenerated amorphous cellulose**<br>Avicel (42%)<br>Hydroxyethylcellulose (13%)<br>CMC (11%)<br>Oat spelts xylan (3%) | n.d. | n.d. | (Zhang et al., 2010) |
| **-** | cellulose 1,4-beta-cellobiosidase (non-reducing end) (91) | *Streptomyces coelicolor A(3)* SCO6546 | 48 | β-1,4 | - | Exo (non-red. end) | cellobiose | 5.0 | **50°C**<br>30°C (70%) | **Avicel**<br>CMC (97%) | 13.3 $s^{-1}$<br>- | 2.7 mM<br>- | (Lee et al., 2018) |

**Table S2** Changes in viable (green-stained), damaged (red- and dual-stained) and unstained oospores in treated oospore samples as determined by live/dead fluorescent staining using fluorescein diacetate/DiTO-3 dyes and microscopic enumeration, repeated measures one-way ANOVA analysis and effect sizes.

| Comparison (1 vs. 2) | Type of oospores | Mean 1 (%)[a] | Mean 2 (%)[a] | Mean diff. (%) [b] | 95% CI of diff. (%) | Adjusted P value | SD of diff.[c] | d[d] | Effect size |
|---|---|---|---|---|---|---|---|---|---|
| Enzyme vs. Buffer | Viable | 43 | 58 | -15 | -57 to 27 | 0.7163 | 16 | 0.9 | Large |
| | Damaged | 41 | 36 | +6 | -41 to 52 | 0.9998 | 18 | 0.3 | Small |
| | Unstained | 15 | 6 | +9 | -42 to 61 | 0.9924 | 20 | 0.5 | Medium |
| SteriGENE vs. Buffer | Viable | 37 | 58 | -21 | -93 to 50 | 0.8574 | 28 | 0.8 | Large |
| | Damaged | 57 | 36 | 21 | -54 to 96 | 0.8858 | 29 | 0.7 | Medium |
| | Unstained | 6 | 6 | 0 | -6 to 6 | >0.9999 | 2 | 0.1 | - |
| Enz. + SteriGENE vs. SteriGENE | Viable | 22 | 37 | -15 | -71 to 41 | 0.9111 | 22 | 0.7 | Medium |
| | Damaged | 74 | 57 | 17 | -34 to 68 | 0.7843 | 20 | 0.9 | Large |
| | Unstained | 4 | 6 | -2 | -16 to 11 | 0.9963 | 5 | 0.4 | Small |
| Mandipropamid vs. 1% DMSO | Viable | 40 | 37 | 4 | -24 to 32 | 0.9993 | 11 | 0.3 | Small |
| | Damaged | 52 | 51 | -1 | -34 to 35 | >0.9999 | 13 | 0.1 | - |
| | Unstained | 7 | 12 | -5 | -16 to 7 | 0.6214 | 5 | 1.0 | Large |
| Enz. + Mandipropamid vs Mandipropamid | Viable | 31 | 40 | -10 | -47 to 28 | 0.9329 | 15 | 0.7 | Medium |
| | Damaged | 64 | 52 | 12 | -19 to 43 | 0.6654 | 12 | 1.0 | Large |
| | Unstained | 5 | 7 | -2 | -10 to 5 | 0.8208 | 3 | 0.9 | Large |
| Copper sulfate vs. Buffer | Viable | 38 | 58 | -21 | -119 to 78 | 0.9782 | 39 | 0.5 | Medium |
| | Damaged | 17 | 36 | -19 | -51 to 14 | 0.3006 | 13 | 1.4 | Very large |
| | Unstained | 45 | 6 | 39 | -32 to 110 | 0.3183 | 28 | 1.4 | Very large |
| Enz. + Copper sulfate vs. Copper suflate | Viable | 26 | 38 | -11 | -44 to 22 | 0.7657 | 13 | 0.9 | Large |
| | Damaged | 28 | 17 | 10 | -43 to 64 | 0.9876 | 21 | 0.5 | Medium |
| | Unstained | 46 | 45 | 0 | -36 to 37 | >0.9999 | 14 | <0.1 | - |
| Metalaxyl-M. vs 1% DMSO | Viable | 42 | 37 | 6 | -11 to 22 | 0.7492 | 6 | 0.9 | Large |
| | Damaged | 51 | 51 | -1 | -33 to 32 | >0.9999 | 13 | 0.1 | - |
| | Unstained | 7 | 12 | -5 | -22 to 13 | 0.9242 | 7 | 0.7 | Medium |
| Enz. + Metalaxyl-M vs. Metalaxyl-M | Viable | 36 | 42 | -7 | -65 to 52 | 0.9999 | 23 | 0.3 | Small |
| | Damaged | 56 | 51 | 6 | -44 to 55 | 0.9998 | 19 | 0.3 | Small |
| | Unstained | 8 | 7 | 1 | -15 to 16 | >0.9999 | 6 | 0.1 | - |
| Nikkomycin Z vs. 1% DMSO | Viable | 50 | 37 | 13 | -9 to 36 | 0.2606 | 9 | 1.5 | Very large |
| | Damaged | 43 | 51 | -8 | -38 to 23 | 0.9344 | 12 | 0.7 | Medium |
| | Unstained | 7 | 12 | -5 | -24 to 13 | 0.8690 | 7 | 0.8 | Large |
| Enz. + Nikkomycin Z vs. Nikkomycin Z | Viable | 65 | 50 | 15 | -35 to 65 | 0.8661 | 20 | 0.8 | Large |
| | Damaged | 30 | 43 | -13 | -66 to 40 | 0.9459 | 21 | 0.6 | Medium |
| | Unstained | 5 | 7 | -2 | -17 to 13 | 0.9995 | 6 | 0.3 | Small |

[a] n = 5
[b] mean of pairwise differences
[c] standard deviation of the pairwise differences, calculated between two treatments for each replicate pair, separate from ANOVA analysis
[d] Cohen's d = mean diff. / SD of diff.
